# Profiling Tel1 Signaling Reveals a Non-Canonical Motif Targeting DNA Repair and Telomere Control Machineries

**DOI:** 10.1101/2024.07.03.601872

**Authors:** Will Comstock, Ethan Sanford, Marcos Navarro, Marcus B Smolka

## Abstract

The stability of the genome relies on Phosphatidyl Inositol 3-Kinase-related Kinases (PIKKs) that sense DNA damage and trigger elaborate downstream signaling responses. In *S. cerevisiae*, the Tel1 kinase (ortholog of human ATM) is activated at DNA double strand breaks (DSBs) and short telomeres. Despite the well-established roles of Tel1 in the control of telomere maintenance, suppression of chromosomal rearrangements, activation of cell cycle checkpoints, and repair of DSBs, the substrates through which Tel1 controls these processes remain incompletely understood. Here we performed an in depth phosphoproteomic screen for Tel1-dependent phosphorylation events. To achieve maximal coverage of the phosphoproteome, we developed a scaled-up approach that accommodates large amounts of protein extracts and chromatographic fractions. Compared to previous reports, we expanded the number of detected Tel1-dependent phosphorylation events by over 10-fold. Surprisingly, in addition to the identification of phosphorylation sites featuring the canonical motif for Tel1 phosphorylation (S/T-Q), the results revealed a novel motif (D/E-S/T) highly prevalent and enriched in the set of Tel1-dependent events. This motif is unique to Tel1 signaling and not shared with the Mec1 kinase, providing clues to how Tel1 plays specialized roles in DNA repair and telomere length control. Overall, these findings define a Tel1-signaling network targeting numerous proteins involved in DNA repair, chromatin regulation, and telomere maintenance that represents a framework for dissecting the molecular mechanisms of Tel1 action.

## Introduction

Phosphatidyl Inositol 3-Kinase-related Kinases (PIKKs) play key roles in genome maintenance by sensing DNA damage and orchestrating DNA damage response signaling networks. In *Saccharomyces cerevisiae,* the PIKKs Mec1 (human ATR) and Tel1 (human ATM) play central roles in coordinating DNA damage responses. While Mec1 senses and responds to single-stranded DNA (ssDNA) via Ddc2 (human ATRIP), Tel1 associates with the blunt ends of double-stranded DNA breaks (DSBs) and short telomeres (1–3) (Figure 1A). Recruitment of either Mec1 or Tel1 to DNA lesions promotes the activation of those kinases, which then initiates a signaling cascade that includes phosphorylation of downstream DNA damage checkpoint kinases Dun1, Rad53, and Chk1 (4–6). Mec1 and Tel1 play partially redundant roles in the orchestration of DNA damage responses. For example, both kinases are important for the suppression of gross chromosomal rearrangements (GCRs), with GCRs accumulating synergistically upon deletion of both kinases (7). Mec1 also plays key roles in surveilling and controlling DNA replication and replication stress responses, most of which are not shared with Tel1 (8). On the other hand, Tel1 plays crucial roles in telomere maintenance by binding to short telomeres during late S/G2 and promoting their elongation (1, 3). Moreover, Tel1 is also involved in meiotic DSB interference (9). The downstream targets through which Tel1 controls these processes and its other related roles in DSB response and repair remain incompletely understood. Tel1 is activated by recruitment to blunt DNA ends via the MRX complex, composed of Mre11, Rad50, and Xrs2 (10). This complex is rapidly recruited to DSBs and plays roles in DNA end-recognition and end-processing. Within the MRX complex, Xrs2 is responsible for interacting with and activating Tel1 via its C-terminal Tel1-interacting region (11).

**Figure 1:**
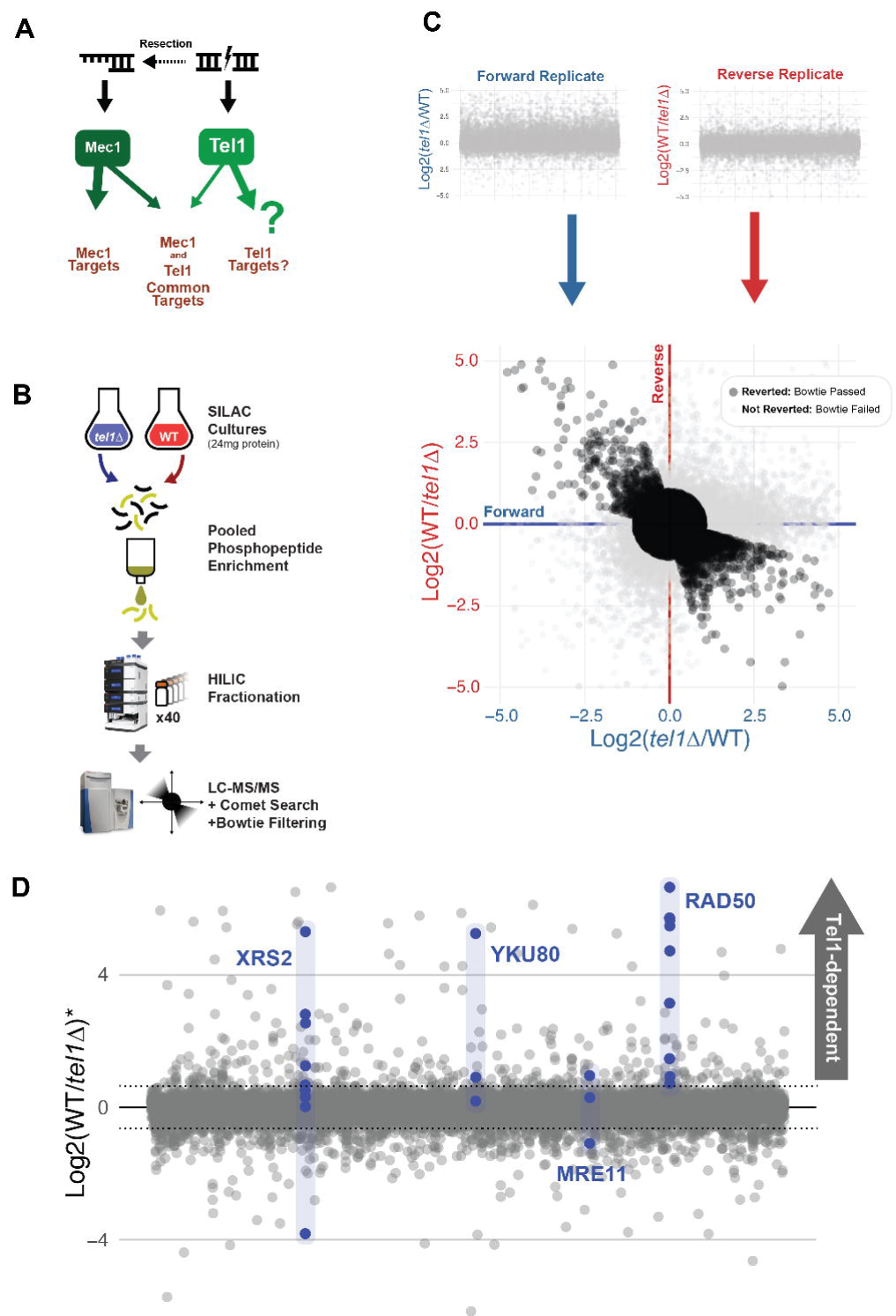
Large scale phosphoproteomics of Tel1-dependent phosphorylation. (A) Overview of DNA lesions activating PIKKs, with Mec1 and Tel1 canonically becoming activated by different forms of DNA damage. (B) Schematic of large-scale phosphoproteomic pipeline. Large SILAC yeast cultures are required to obtain sufficient starting material for tryptic digestion and phosphopeptide enrichment. Extensive HILIC prefractionation reduces sample complexity and allows for in-depth LC-MS/MS analysis, followed by database searching and Bowtie replicate filtering. (C) Bowtie filtering of forward and reverse replicates performed for each experimental condition, in which experimental conditions are inverted between SILAC channels. The replicates are then plotted against one another to determine which phosphopeptides feature appropriately inverting quantitative ratios, discarding any that do not invert. (D) The quantitative ratios averaged between forward and reverse replicates for 17,486 phosphopeptides that passed Bowtie filtering. Among these are phosphopeptides on Yku80 and components of the MRX complex, most of which are found to be Tel1-dependent.

Mec1 and Tel1 are well known to preferentially phosphorylate motifs featuring serine or threonine residues followed by a glutamine residue (S/T-Q motif) (2). Several phosphoproteomic screens have been performed to map the signaling network mediated by Mec1, which led to the identification of hundreds of putative Mec1 targets phosphorylated at the S/T-Q motif (12–14). Comparatively, a much smaller set of Tel1 targets were identified, consistent with the notion that Tel1 contributes less to the DNA damage signaling response overall relative to Mec1. Our previous phosphoproteomic screen for Tel1 targets identified only 4 highly Tel1-dependent phosphorylation events, including Rad50, Enp1, Asg1, and Tfg1 (14). Other known Tel1 targets include Cdc13 and Rad53, and Pol4 (15, 16).

In an effort to map the Tel1 signaling network with higher coverage, we deployed a scaled-up phosphoproteomic pipeline alongside our previously published “bowtie” filtering methodology (17). *tel1*Δ yeast strains were compared against wild type yeast using stable isotope labeling by amino acids in cell culture (SILAC) mass spectrometry, from which hundreds of novel Tel1-dependent sites were detected. As expected, a significant proportion of these sites featured the canonical S/T-Q motif. Surprisingly, we also detected a strong prevalence and enrichment of a novel acidophilic consensus motif (D/E-S/T) among Tel1-dependent phosphorylation events. Overall, our results define a Tel1-signaling network targeting proteins involved in DNA repair, chromatin, and telomere maintenance that provides a framework for dissecting the molecular mechanisms of Tel1 action.

## Materials and Methods

### Yeast Strains and Culture

A complete list of yeast strains used in this study can be found in supplemental table 2. ORF deletions were performed via the established polymerase chain reaction (PCR)-based strategy to amplify resistance cassettes with flanking sequence homologous to a target gene (Longtine et al 1998). All ORF deletions were verified by PCR) with one set of primers targeting the wild-type sequence and one set of primers targeting the null DNA sequence. Yeast strains were grown at 30°C in a shaker incubator set to 220 rpm. For strains with integrated genetic modifications, YEP-D media was used. For SILAC cultures, yeast strains were grown in -Arg -Lys media supplemented with either isotopically normal arginine and lysine or the ^13^C^15^N isotopologue. Excess proline to prevent arginine to proline conversion was added to SILAC media at a concentration of 80 mg/L.

### CRISPR Mutation

The Hrr25 analog-sensitive mutant was created by first ligating duplexed primers containing the sequence for gRNA targeting sequences corresponding to Hrr25 Isoleucine 82 into a CRISPR-Cas9 vector. Candidate gRNA sequences were identified using CHOPCHOP (18–20). This plasmid was transformed into yeast along with a gBlock DNA segment containing the corresponding Ile82Gly substitution, after which transformants were plated on -Leu to select for yeast expressing the plasmid. After two days, colonies were selected and submitted for Sanger sequencing to determine whether the mutation occurred. Positive clones were then cultured and plated sparsely on YPD agar before being stamped on -Leu to select for clones that had lost the CRISPR-Cas9 plasmid.

### Spot Assays

For dilution assays, 5 mL of yeast culture was grown to saturation at 30°C. One OD600 equivalent of the saturated culture was then 5-fold serially diluted in a 96-well plate in sterile water. Dilutions were then spotted onto agar plates using a bolt pinner.

### Phosphoproteomics

For phosphoproteomics experiments, 300 mL SILAC cultures were grown in heavy or light SILAC media to mid-log phase and treated as described in the figure legends.

Cells were pelleted at 1000 rcf and washed with TE buffer containing 1mM PMSF. Pellets were lysed by bead beating with 0.5 mm glass beads for three cycles of 10 minutes with 1 minute rest time between cycles at 4°C in lysis buffer (150 mM NaCl, 50 mM Tris pH 8.0, 5 mM EDTA, 0.2% Tergitol type NP-40) supplemented with complete EDTA-free protease inhibitor cocktail (Pierce), 5 mM sodium fluoride, and 10 mM D-glycerophosphate. 12 mg of each light- and heavy-labeled protein lysate was pooled, and the pooled lysates were then denatured and reduced with 1% SDS and 5 mM DTT at 42°C, and then alkylated with 25 mM iodoacetamide. Lysates were mixed with cold PPT solution (49.9% EtOH, 50% acetone, 0.1% acetic acid) to precipitate on ice for 30 minutes, after which precipitated protein was pelleted via centrifugation. Pellets were washed once with PPT and then resuspended in Urea/Tris solution (8M urea, 50 mM Tris pH 8.0). Urea-solubilized pellet was then diluted to 1M urea using NaCl/Tris solution (150mM NaCl, 50 mM Tris pH 8.0) and digested overnight at 37°C with TPCK-treated trypsin. Phosphoenrichment for each replicate was performed using two Thermo Fisher High Select Fe-NTA phosphopeptide enrichment kits (cat# A32992) according to the manufacturer’s protocol. Purified phosphopeptides were then extensively fractionated using HILIC chromatography on an Ultimate 3000 HPLC (40 fractions collected), after which fractions were dried via vacuum concentration, resuspended in 0.1% TFA, and subjected to LC-MS/MS analysis on a Thermo Fisher Q-Exactive HF mass spectrometer using 70 minute reverse-phase C18 gradients.

### Mass Spectrometry Data Analysis

Raw MS/MS spectra were searched using the Comet search engine (part of the Trans Proteomic Pipeline; Seattle Proteome Center) over a composite yeast protein database consisting of both the normal yeast protein sequences downloaded for the Saccharomyces Genome Database (SGD) and their reversed protein sequences to serve as decoys and estimate the false discovery rate (FDR) in search results (21).

Search parameters allowed for semi-tryptic peptide ends, a mass accuracy of 15 ppm for precursor ions, variable modifications for SILAC lysine and arginine (8.0142 and 10.00827 daltons, respectively), variable modification for STY phosphorylation (79.966331 daltons), and a static mass modification of 57.021465 daltons for alkylated cysteine residues. Phosphorylation site localization probabilities were determined using PTMProphet, with SILAC quantification of identified phosphopeptides being performed using XPRESS (both modules part of the Trans Proteomic Pipeline; Seattle Proteome Center). Tel1 SILAC phosphoproteomic data was subject to Bowtie filtering as previously described in Faca et al. 2020. Processed phosphoproteomic data is available in supplemental table 2.

### Data Availability

All mass spectrometric data generated for this study are available through PRIDE via PXD identifier PXD053305 (https://www.ebi.ac.uk/pride/).

### Protein Structure Analysis

Structures for Tel1 (PDB ID 5YZ0) and Mec1 (PDB ID 7Z87) were aligned in PyMol while superimposing a p53 peptide substrate from a structure of human ATM bound to this peptide (PDB ID 8OXO).

## Results

### Large scale phosphoproteomic analysis of Tel1-dependent signaling

Our previous phosphoproteomic screen of the Mec1 and Tel1 signaling networks in budding yeast resulted in the identification of a small set of Tel1-dependent phosphorylation events (14). In order to map the Tel1-signaling network with greater depth, we employed a scaled-up phosphoproteomic pipeline (Figure 1B). This pipeline involved scaling up yeast cultures to obtain at least 12 milligrams of extracted protein per experimental condition, followed by multiple IMAC phosphopeptide enrichments in parallel and extensive sample prefractionation (over 30 fractions) via hydrophilic interaction liquid chromatography (HILIC) prior to LC-MS/MS analysis. Yeast cultures were grown in SILAC media for incorporation of light or heavy amino acids that were used for relative quantification. To increase confidence in the identification, quantification and phosphorylation site assignment, we used a Bowtie filtering method in which the SILAC channels were reversed (“forward” and “reverse” replicates) in the second replicate of every phosphoproteomic analysis (17). Cultures of wild type and *tel1*Δ cells were treated with 0.1% methyl methanesulfonate (MMS) for 90 minutes to induce DNA double-stranded DNA breaks (DSBs). Experiments were performed swapping heavy and light amino acids for each genotype, after which these forward and reverse replicates were plotted against one another and their ratios averaged (Figure 1C). In total, 23,166 phosphopeptides were detected, of which 17,486 passed the bow tie filtering and 439 phosphorylation sites and clusters were found to be Tel1-dependent (with at least 50% reduction in abundance in *tel1*Δ cells compared to wild type) (Figure 1D). As expected, components of the MRX complex and DSB repair protein YKU80 were found to contain several Tel1-dependent phosphorylation sites, validating our dataset (Figures 1D, S1).

### Tel1-dependent phosphorylation is enriched in S/T-Q motifs on proteins involved in chromatin and chromosome biology

Consistent with the known preferred motif for Tel1 phosphorylation, there was an enrichment for S/T-Q motif among the set of Tel1-dependent sites uncovered. 16.5% of Tel1-dependent sites were found to feature this motif compared to 5.6% of the group of phosphorylation sites not affected by *TEL1* deletion (Figure 2A). With the identification of 58 Tel1-dependent phosphorylation sites featuring this S/T-Q motif (63 including clusters), the number of Tel1-dependent S/T-Q sites has been expanded 14-fold over our previous phosphoproteomic screen for Tel1 targets (Figure 2B). GO term analysis revealed an enrichment in proteins involved in chromatin and chromosome organization (Figure 2C) (22). Proteins featuring Tel1-dependent phosphorylation at S/T-Q sites also included proteins with established roles in DNA repair and telomere maintenance, such as MSH6, RIF1, YKU80, and RAD50 (Figures 2A and 2D), and represent potential mechanistic links between Tel1 and its roles in controlling telomere length. Tel1-dependent phosphorylation events at the S/T-Q motif were also present on proteins implicated in transcription, indicating potential roles for canonical Tel1 signaling in transcriptional regulation (Figure S2) (23).

**Figure 2:**
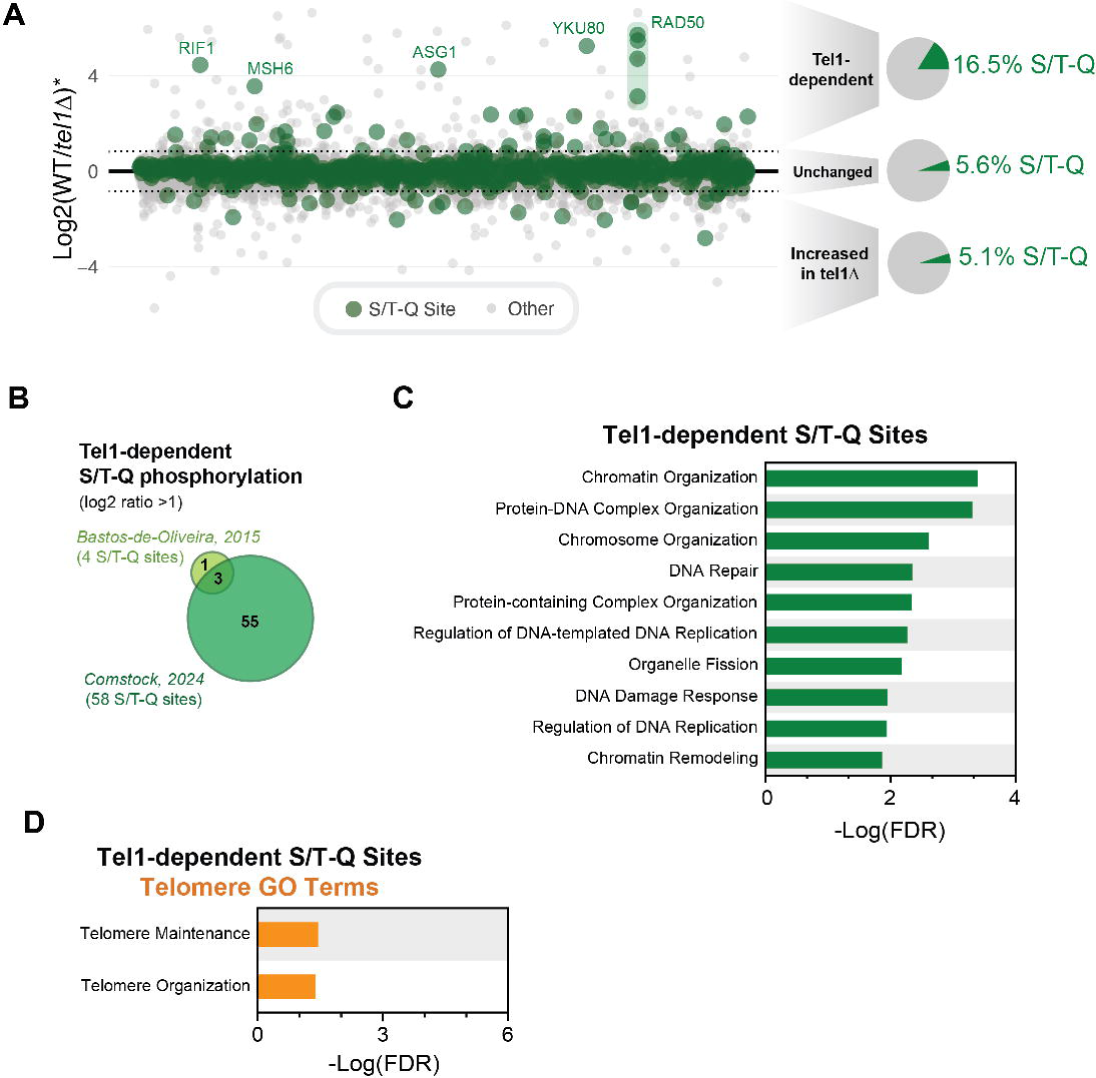
Tel1-dependent phosphorylation events are enriched for the S/T-Q motif on proteins involved in the DNA damage response and telomere maintenance. (A) The averaged quantitative ratio for 17,486 phosphopeptides that passed Bowtie filtering with S/T-Q sites highlighted in green. Tel1-dependent phosphorylation events are over 3X enriched for the canonical preferred PIKK S/T-Q motif. (B) Comparison of the number of Tel1-dependent phosphorylation events detected in the current study versus our previous investigation of the Tel1 signaling network. (C) The top 10 GO terms (minus terms containing “process”) enriched among proteins featuring Tel1-dependent S/T-Q sites. (D) Telomere-related GO term enrichment among proteins featuring Tel1-dependent S/T-Q sites.

### A novel D/E-S/T motif is enriched among Tel1-dependent phosphorylation events

Unexpectedly, amino acid residue prevalence analysis of the Tel1-dependent sites revealed a strong enrichment for an acidic residue at the −1 position relative to the phosphorylation site (Figure 3A). Prevalence analysis revealed that the D/E-S/T motif represented nearly 50% of the Tel1-dependent sites identified, a higher proportion compared to the ∼15% comprised of the S/T-Q motif (Figure 3B-C). Notably, S/T-Q sites preceded by an acidic residue (D/E-S/T-Q motif) were drastically enriched in the set of Tel1-dependent sites, displaying an almost 20-fold increase in prevalence compared to the set of sites not altered by *TEL1* deletion, suggesting a strong preference for this motif by Tel1 (Figure 3C). The set of Tel1-dependent phosphorylation at the D/E-S/T motif was strongly enriched for proteins involved in DNA damage response and DNArepair and contained more GO terms related to telomere biology compared to the set of S/T-Q motifs (Figures 3D-E). These results show that the proteins with phosphorylation at the D/E-S/T motif tend to align with known Tel1 functions and suggest that this novel Tel1 signaling motif reflects phosphorylation events through which Tel1 exerts its regulatory roles. Proteins implicated in RNA metabolic processes were also prevalent among the 189 Tel1-dependent D/E-S/T phosphorylation sites and clusters, including a prominent connected node of proteins implicated in ribosomal RNA processing (Figure S3).

**Figure 3:**
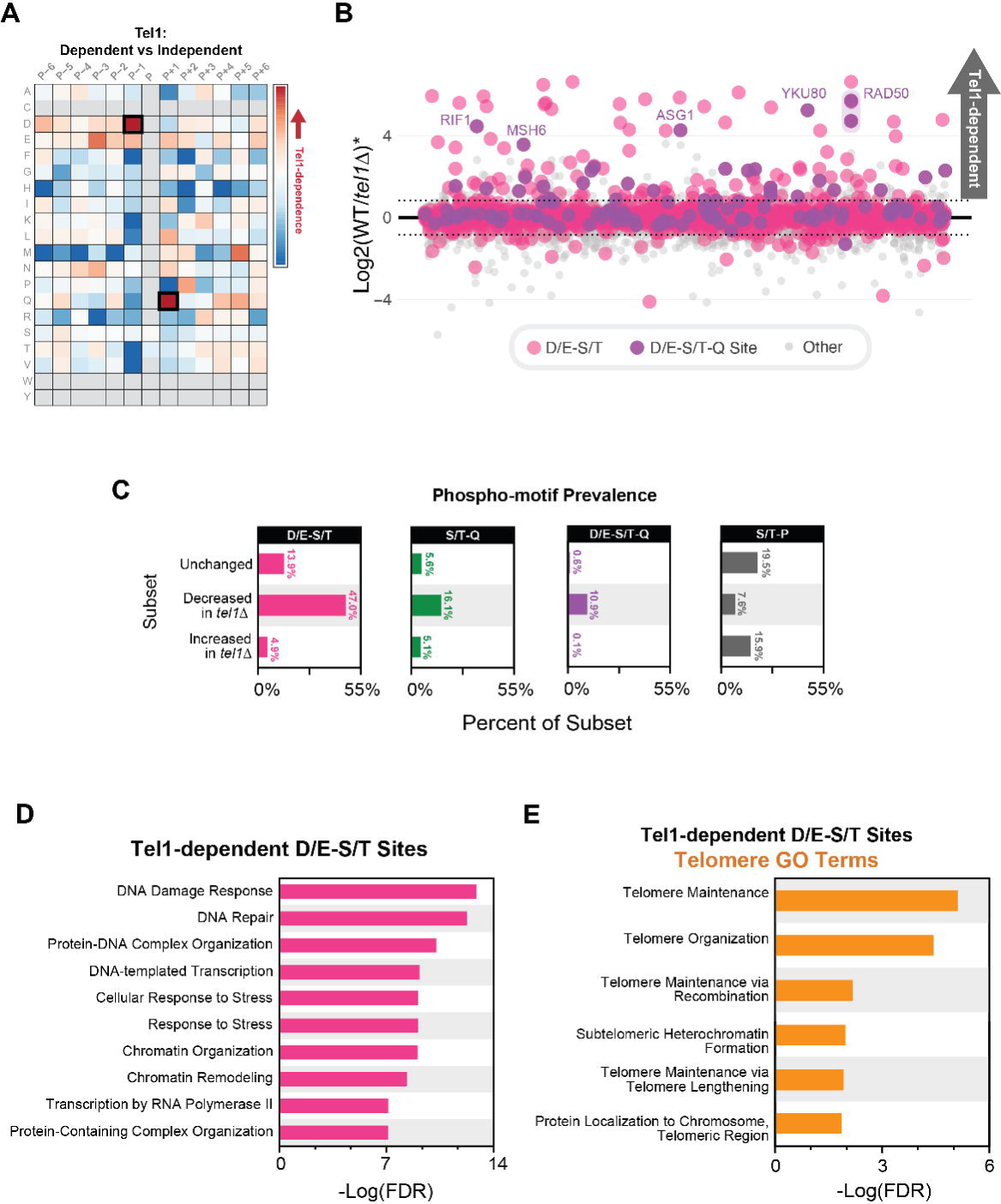
A novel D/E-S/T motif is more enriched than the S/T-Q motif among Tel1-dependent phosphorylation events. (A) Heatmap illustrating enrichment of amino acid residues at loci relative to phosphorylation events found to be Tel1-dependent. Acidic residues, particularly aspartic acid, are found to be enriched in the −1 locus. Enrichment for the S/T-Q motif is also illustrated here. (B) The averaged quantitative ratio for 17,486 phosphopeptides that passed Bowtie filtering with D/E-S/T sites highlighted in pink and the sites with the combined motif (D/E-S/T-Q) highlighted in purple. (C) Prevalence of four phospho-motifs among subsets of the Tel1 signaling dataset. The most enriched motif in phosphopeptides that decreased upon deletion of Tel1 was D/E-S/T, with S/T-Q and D/E-S/T-Q motifs also showing high enrichment among this same subset of the data. (D) The top 10 GO terms (minus terms containing “process”) enriched among proteins featuring Tel1-dependent D/E-S/T sites. (E) Telomere-related GO term enrichment among proteins featuring Tel1-dependent D/E-S/T sites.

### Mec1-dependent phosphorylation events do not feature D/E-S/T motif enrichment

We next asked if enrichment of the D/E-S/T consensus motif is also observed in Mec1-dependent phosphorylation events or if it is a feature specific to Tel1-dependent signaling. We compared data from our previous study mapping Mec1-dependent signaling to the Tel1-dependent dataset identified here (Figure 4A) (24). Interestingly, no enrichment for the D/E-S/T consensus motif was found among Mec1-dependent phosphorylation events, whereas the canonical S/T-Q phosphorylation motif was more strongly enriched among Mec1-dependent phosphorylation sites than it was among Tel1-dependent sites (Figure 4B, C). This difference aligns with existing structural data showing that the Mec1 pocket for recognition of substrate residues displays a negatively charged distribution at the −1 position, primarily due to the presence of an E^2228^ in its catalytic loop (Figure 4D). In contrast, Tel1 features an asparagine (N2616) at this position, which, together with basic residues R^2544^ and H^2526^, creates a more positively charged distribution that favors acidic residues at the −1 position of substrates (Figure 4D) (25–27). These findings reveal that the phosphorylation of D/E-S/T is a feature specific to Tel1 signaling. Interestingly, we observed that several Tel1 signaling events mapped here are induced in cells lacking Mec1 (lower-right quadrant in Figure 4A), consistent with genetic data showing that in the absence of Mec1, Tel1 plays an important compensatory role in preventing GCRs, slow growth, and DNA damage sensitivity (8, 28).

**Figure 4:**
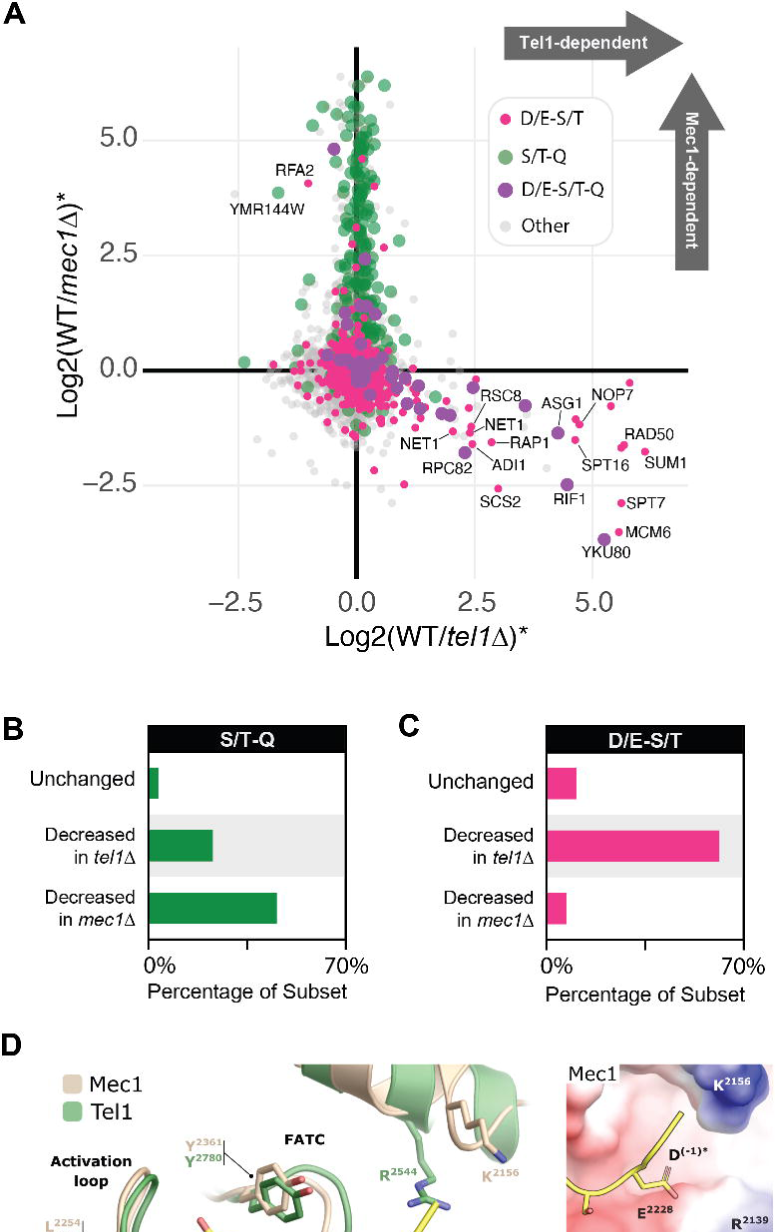
The D/E-S/T motif is more enriched in Tel1-dependent phosphorylation events compared to Mec1-dependent phosphorylation events. (A) Tel1 phosphoproteomic data from the current study plotted against Mec1 phosphoproteomic data from Sanford et al. 2021 in which both Mec1 and adaptor protein Rad9 were deleted to investigate Mec1-dependent signaling events. Increased abundance of Tel1-dependent phosphorylation events upon loss of Mec1 supports the notion that Tel1 engages in some amount of compensatory signaling when Mec1 activity is compromised. (B) Enrichment of the S/T-Q motif among Tel1- and Mec1-dependent phosphorylation events illustrates a higher enrichment for this motif among Mec1-dependent phosphorylation events. (C) Enrichment of the D/E-S/T motif among Tel1- and Mec1-dependent phosphorylation events strongly suggests that D/E-S/T motif enrichment is exclusive to Tel1-dependent signaling. (D) Structural superimposition of yeast Mec1 and Tel1 (PDB IDs 7Z87 and 5YZ0 respectively) highlighting the substrate-binding pocket and key residues involved in the recognition of peptide substrates via their positions −1 (Tel1 H^2526^, R^2544^, and N^2616^; Mec1 R^2139^, K^2156^ and E^2228^) and +1 (Tel1 L^2642^ and Y^2780^; Mec1 L^2254^ and Y^2361^). Inserts on the right show the electrostatic potential of surfaces around the −1 binding pocket for each kinase, blue indicating positive charge and red indicating negative charge. In all figures, the peptide substrate shown in yellow is superimposed from a structure of ATM bound to p53 (PDB ID 8OXO) with the original Leu at the −1 position remodeled as Asp (asterisk). ATP loops of kinases are omitted to facilitate visualization. Carbon atoms are tan for Mec1 and green for Tel1.

### Tel1-dependent phosphorylation events at the D/E-S/T motif are not dependent on downstream kinases Dun1, Rad53, or Hrr25

The Tel1-dependent phosphorylation events featuring the D/E-S/T motif may represent direct substrates of Tel1 or could be targeted by other downstream kinases under the control of Tel1. To determine the effect of downstream kinases on the phosphorylation status of Tel1-dependent phosphorylation events at the D/E-S/T motif, we searched for kinases containing Tel1-dependent phosphorylation in our dataset, which would point to kinases potentially activated by Tel1. This analysis revealed that the Hrr25 and Dun1 kinases are potential downstream kinases under the control of Tel1-dependent signaling (Figure 5A). Consistent with this notion, Hrr25 and Dun1 contain a Tel1-dependent S/T-Q phosphorylation site at their regulatory and kinase domains, respectively (Figure 5B-C). We next performed phosphoproteomic analyses in cells defective for these kinases to monitor the effect on Tel1-dependent phosphorylation events at the D/E-S/T motif.

**Figure 5:**
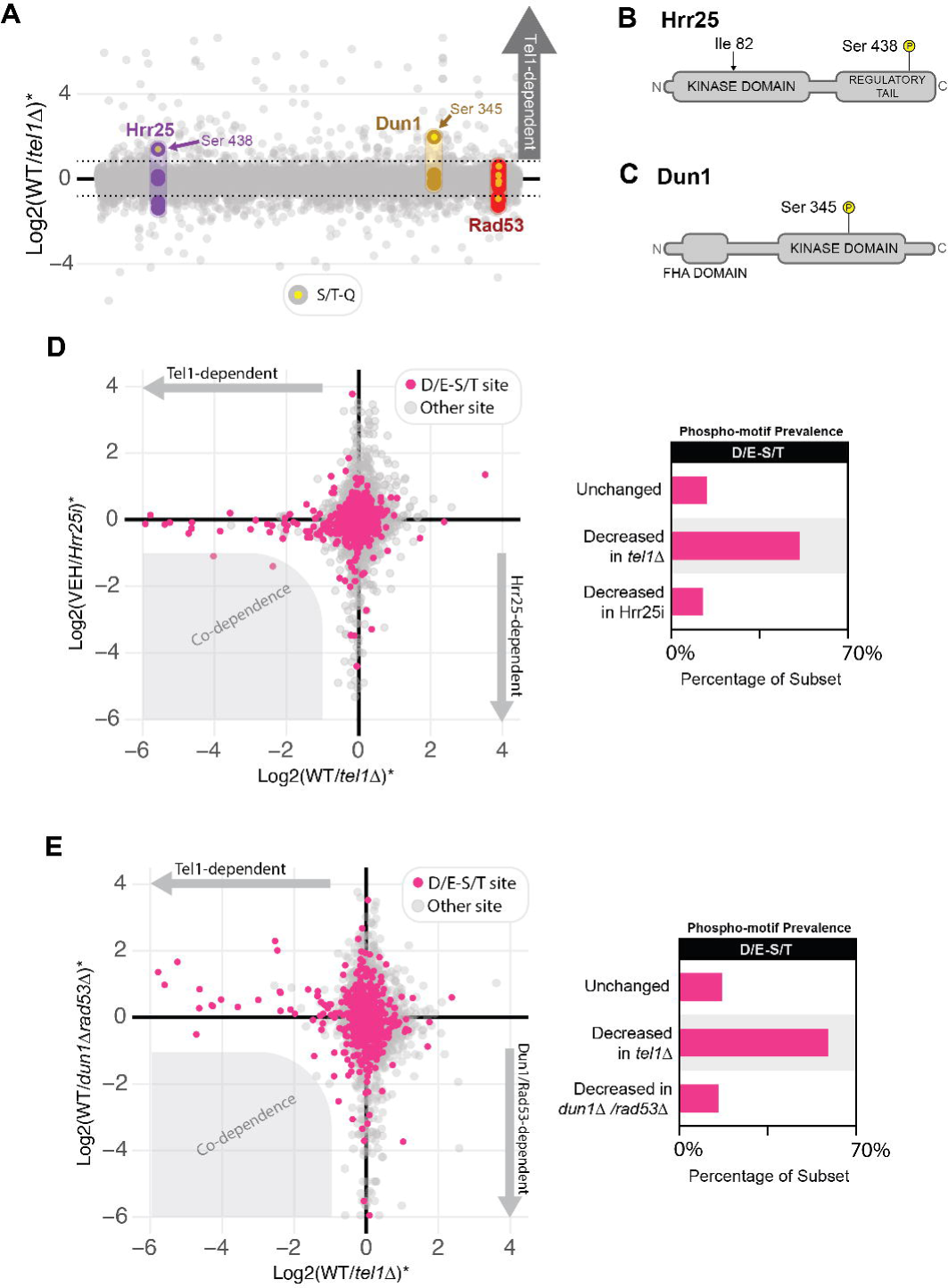
Tel1-dependent D/E-S/T phosphorylation events are not dependent on downstream kinases Hrr25, Dun1, or Rad53. (A) S/T-Q sites on kinases Hrr25 and Dun1 were found to be Tel1-dependent. (B) Sequence map of Hrr25 showing where an analog-sensitivity mutation was made (Ile 82) as well as the location of the Tel1-dependent S/T-Q phosphorylation event (Ser 438). (C) Sequence map of Dun1 showing the location of the Tel1-dependent S/T-Q phosphorylation event (Ser 345). (D) Tel1 phosphoproteomic data plotted against Hrr25 phosphoproteomic data generated using acute kinase inhibition via analog sensitivity mutation, with no significant co-dependence of signaling events observed. The D/E-S/T motif was not found to be enriched among Hrr25-dependent phosphorylation events. (E) Tel1 phosphoproteomic data plotted against Dun1/Rad53 phosphoproteomic data generated using deletion of both kinases, with no significant co-dependence of signaling events observed. The D/E-S/T motif was not found to be enriched among Dun1/Rad53-dependent phosphorylation events.

Given that Hrr25 is essential for cell survival, a documented ATP analog-sensitive mutation was created (Hrr25as) to allow for screening of Hrr25-dependent phosphorylation events (29). This system enables acute inhibition of Hrr25, which was confirmed by the observed increased cell sensitivity to both the ATP analog (1NM-PP1) and MMS in the presence of the analog (Figure S4). Phosphoproteomic analysis revealed no enrichment for the D/E-S/T consensus motif in the set of Hrr25-dependent phosphorylation events (Figure 5D). Importantly, none of the Tel1-dependent phosphorylation events at the D/E-S/T motif were found to be strongly reduced upon impairment of Hrr25 (no co-dependence). These results rule out Hrr25 as the kinase responsible for phosphorylating the Tel1-dependent phosphorylation events at the D/E-S/T motif.

We next tested if Dun1 was required for the phosphorylation of the Tel1-dependent phosphorylation events at the D/E-S/T motif. Notably, we did not detect Tel1-dependent phosphorylation on Rad53 (Figure 5A), a kinase involved in the canonical activation of Dun1, consistent with previous reports suggesting that Dun1 might also be directly activated by Tel1 (6). Nonetheless, to rule out any contribution from Rad53 and/or Dun1, we utilized a cell lacking both kinases in the phosphoproteomic analysis. As with Hrr25, no enrichment for the D/E-S/T consensus motif was observed among Dun1/Rad53-dependent phosphorylation events (Figure 5E). Interestingly, several Tel1-dependent events at the D/E-S/T motif were observed to be up-regulated upon double deletion of *DUN1* and *RAD53* (Figure 5E), suggesting that in the absence of these kinases, Tel1 plays an important compensatory role in the DNA response, similar to what was observed in the absence of Mec1. Overall, these findings rule out the involvement of Dun1, Rad53 and Hrr25 as kinases downstream of Tel1 signaling responsible for the phosphorylation of Tel1-dependent phosphorylation events at the D/E-S/T motif. Overall, these results strengthen the possibility that these sites are directly phosphorylated by Tel1.

### A network of Tel1-dependent phosphorylation connects Tel1 to DNA damage responses and telomere maintenance

Overall, our phosphoproteomic analysis identified 353 phosphorylation sites dependent on Tel1, of which about 53% are located at a D/E-ST, D/E-S/T-Q or S/T-Q motif (Figure 6A). Among proteins featuring these sites, some of the most enriched cellular processes are DNA damage response, transcription, and telomere organization. The majority of the proteins in these processes were phosphorylated at the D/E-S/T or D/E-S/T-Q motif, and have not previously been reported to be regulated by Tel1 phosphorylation (Figure 6B). Given the established role of Tel1 in controlling telomere length, for which the functional Tel1 substrate(s) involved remain(s) unknown, we inspected Tel1-dependent phosphorylation in the set of proteins important for telomere maintenance (Figure 6C).

**Figure 6:**
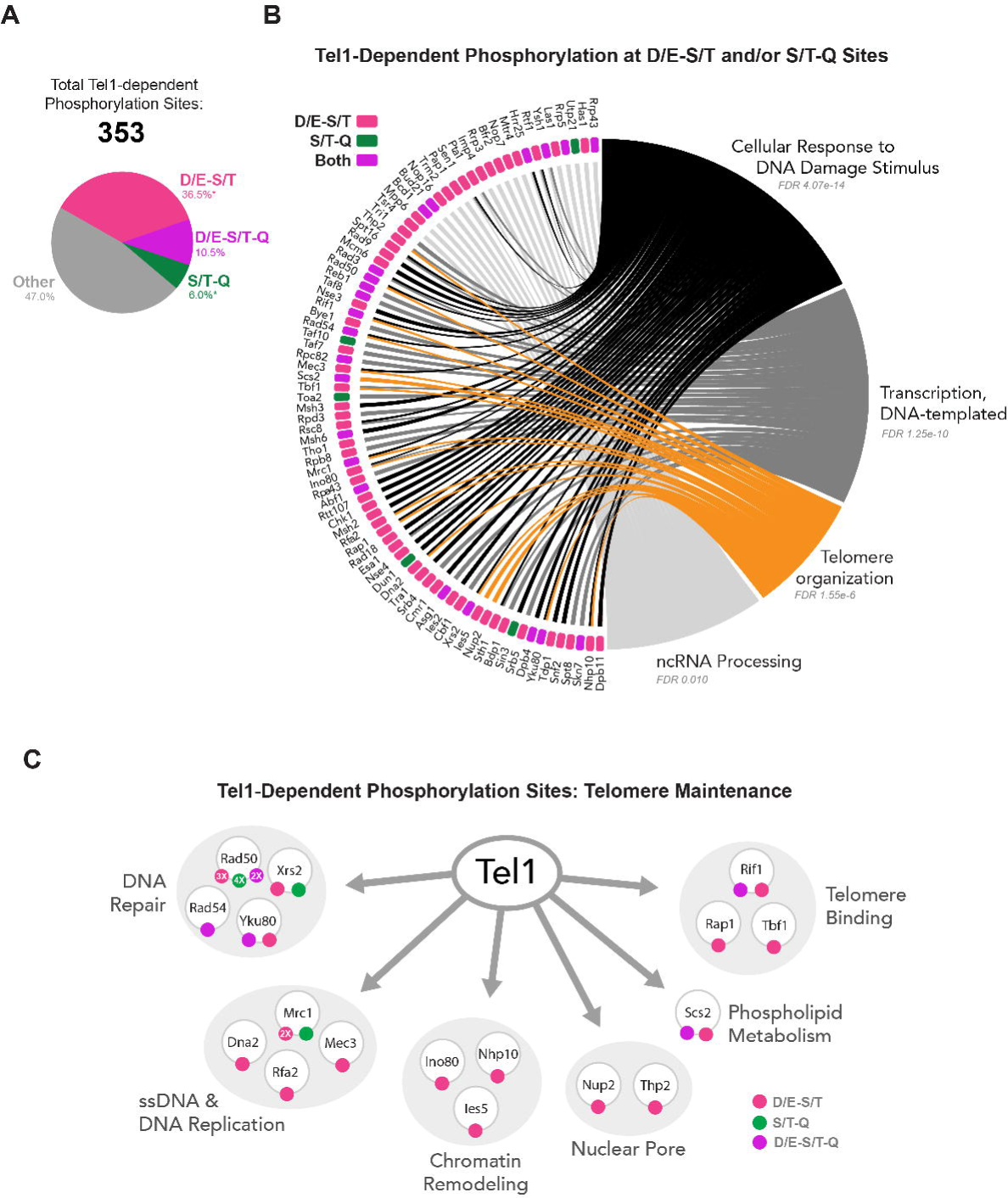
An expanded network of Tel1-dependent phosphorylation targeting S/T-Q and D/E-S/T motifs. (A) Overview of Tel1-dependent phosphorylation events that were able to be localized and their motifs. *D/E-S/T and S/T-Q motif prevalence calculated excluding the combined motif, D/E-S/T-Q. (B) Tel1-dependent phosphorylation events with either the S/T-Q or D/E-S/T motif implicated in the DNA damage response, DNA-templated transcription, telomere organization, or noncoding RNA processing. Proteins denoted as having “both” motifs either contain multiple sites with both motifs represented or at least one site with the combined motif, D/E-S/T-Q. (C) Breakdown of phosphorylation events implicated in telomere organization as denoted by panel B.

The result defines a cohesive network of proteins involved in DNA repair, including components of the MRX complex, as well as Rad54 and YKU80. Three telomere binding proteins intrinsically involved in telomere homeostasis also displayed Tel1-dependent phosphorylation, and represent likely effectors of Tel1 regulation at telomeres. Chromatin remodeling proteins linked to the Ino80 remodeller, which were previously reported to be involved in telomere maintenance, also had Tel1-dependent phosphorylation (30). Other categories included proteins involved in DNA replication, ssDNA transactions, and nuclear pores. Of importance, in the majority of the proteins involved in telomere maintenance found to have Tel1-dependent phosphorylation, the detected phosphorylation site resides at a D/E-S/T motif. These findings provide a framework of proteins and phosphorylation sites to guide functional and mechanistic analyses of Tel1 function, and highlight the importance of this novel motif to expand our knowledge of the network of Tel1 signaling.

## Discussion

Tel1 has long been recognized as a key regulator of telomeres and DSB responses in budding yeast. The molecular mechanisms and specific substrates through which Tel1 controls these processes remains elusive. Here we mapped over 300 phosphorylation events dependent on Tel1, most of which are independent of the downstream kinases Rad53 and Dun1, and represent potential direct targets of Tel1. Consistent with these events being directly mediated by Tel1, we observed an enrichment in the canonical S/T-Q motif for PIKK phosphorylation. Surprisingly, we observed an even greater enrichment for a D/E-S/T motif, which we propose is a novel preferential motif for Tel1 phosphorylation. The identification of this motif and potential novel substrates and sites of Tel1 phosphorylation should provide a more comprehensive framework of the Tel1 signaling network for functional studies aiming at defining the molecular mechanism of Tel1 action.

Several lines of evidence are consistent with the D/E-S/T motif being directly targeted by Tel1. First, the D/E-S/T motif was the most enriched motif in the set of Tel1-dependent phosphorylation and was independent of known downstream kinases (and independent of any other kinase found to carry a Tel1-dependent phosphorylation).

Second, we observed that the D/E-S/T-Q motif, which matches the canonical S/T-Q as well as the novel D/E-S/T motif, was the most enriched motif in the set of Tel1-dependent phosphorylation, occurring at a frequency over 19-fold higher than in the set of Tel1-independent events. Third, analysis of the substrate binding pocket of Tel1 revealed that it contains a positively charged distribution that favors acidic residues at the −1 position of substrates. Fourth, the D/E-S/T preference is specific to Tel1 and absent from signaling mediated by Mec1, whose substrate binding pocket has a negatively charged distribution that does not favor acidic residues at the −1 position of substrates. We note, however, that we cannot fully eliminate the possibility that the identified D/E-S/T motif is being caused indirectly by another kinase downstream of Tel1. Since we were unable to obtain active recombinant Tel1 preparations that were devoid of contaminating kinase activity, we could not confirm the direct phosphorylation of D/E-S/T motif using *in vitro* kinase reactions.

The finding of a non-canonical mode of Tel1 signaling will help understand how Tel1 plays specific roles in telomere maintenance and DNA repair that are not shared with Mec1. Since Tel1 and Mec1 have been reported to share similar substrate specificity toward S/T-Q, and redundantly target many of the same proteins (12, 13), it has been difficult to determine the basis of Tel1’s unique biological functions. We propose that the specificity of Tel1 signaling in targeting D/E-S/T motifs helps explain the specialized roles for Tel1 in the control telomere length and other biological processes.

The ability of PIKKs to target non-S/T-Q motifs has been recently documented in human cells. For example, we recently found that DNA-PKcs can directly phosphorylate an S/T-bulky-D/E motif (31). Additionally, ATR was previously found to auto-phosphorylate at an S/T-P motif (32). While we have not observed an enrichment of D/E-S/T motif in a set of ATM-dependent events we recently reported, we do not exclude the possibility that ATM may also target such a motif, or other non-S/T-Q motifs (31). More in-depth analysis of ATM-dependent signaling in mammalian cells will be needed to properly probe alternative motifs.

## Acknowledgements

We thank Beatriz S. Almeida for technical support and members of the Smolka Lab for valuable discussions.

---

**Figure S1:**
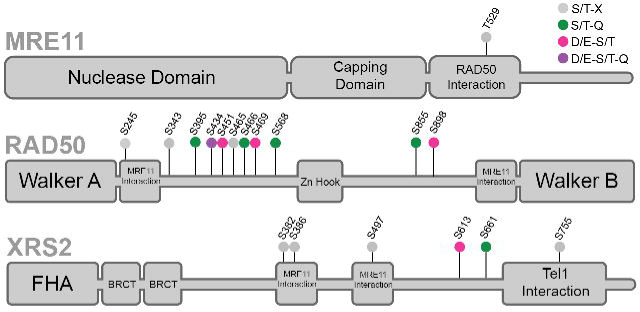
Tel1-dependent phosphorylation sites on components of the MRX complex. Many sites featuring various motifs were found to be Tel1-dependent, with Rad50 featuring the most Tel1-dependent phosphorylation events of any protein in the dataset.

**Figure S2:**
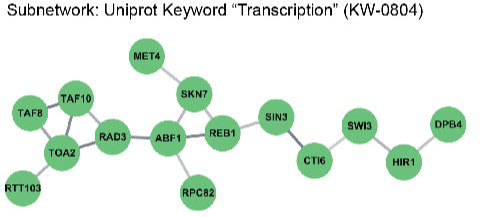
Tel1-dependent S/T-Q phosphorylation events featured on proteins involved in transcription. A subnetwork of proteins annotated with Uniprot keywork “Transcription” was found via STRING network analysis among Tel1-dependent S/T-Q phosphorylation events.

**Figure S3:**
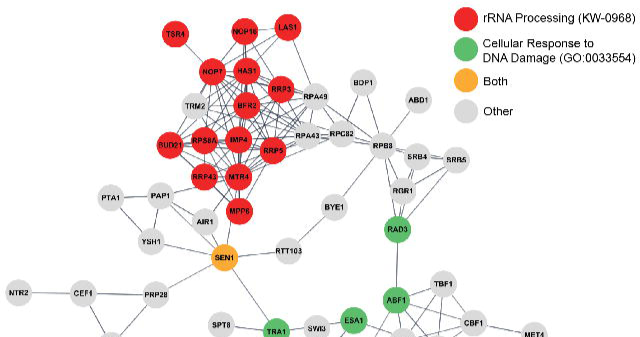
Tel1-dependent D/E-S/T phosphorylation event “RNA Metabolic Process” subnetwork. Many of these phosphorylation events were found to be on proteins involved in RNA processing in addition to the DNA damage response, with a prominent cluster of proteins involved with ribosomal RNA processing specifically.

**Figure S4:**
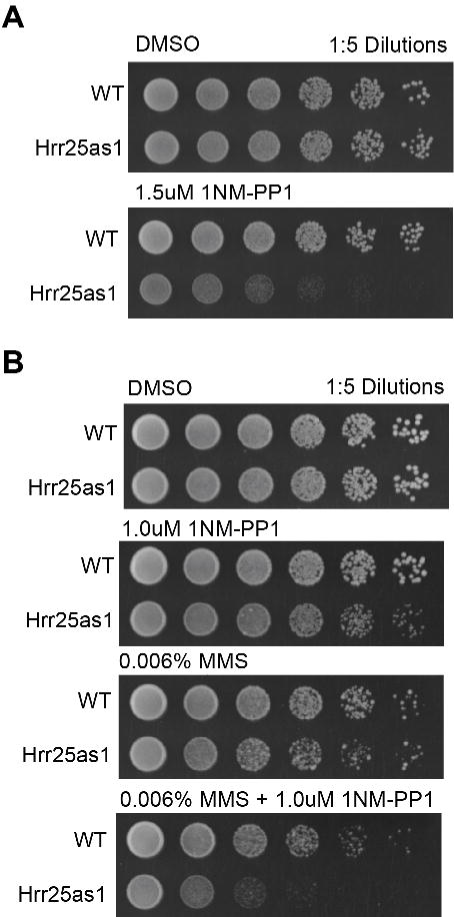
Hrr25 Analog-sensitive mutant (Hrr25as1) shows sensitivity to ATP analog 1NM-PP1 and increased sensitivity to genotoxin MMS. (A) Upon addition of 1.5uM 1NM-PP1 to YPD agar, an analog-sensitive mutant of essential kinase Hrr25 shows attenuated growth. (B) Hrr25as1 becomes sensitive to 0.006% MMS upon addition of a low dose of 1NM-PP1 (1.0uM).

